# Polycomb-mediated transgenerational epigenetic inheritance of Drosophila eye colour is independent of small RNAs

**DOI:** 10.1101/2024.09.26.615209

**Authors:** Maximilian H. Fitz-James, Poppy Sparrow, Christopher Paton, Peter Sarkies

## Abstract

Transgenerational epigenetic inheritance (TEI) describes the process where distinct epigenetic states may be transmitted between generations, resulting in stable gene expression and phenotypic differences between individuals that persist independently of DNA sequence variation. Chromatin modifications have been demonstrated as important in TEI, however, the extent to which they require other signals to establish and maintain epigenetic states is still unclear. Here we investigate whether small non-coding RNAs contribute to different epigenetic states of the Fab2L transgene in *Drosophila* triggered by transient chromatin contacts, which requires Polycomb complex activity to deposit the H3K27me3 modification for long-term TEI. Using mutants deficient in known small non-coding RNAs, high throughput sequencing, investigation of chromatin conformation and gene expression analysis we demonstrate that small non-coding RNAs do not contribute directly to initiation or maintenance of silencing. However, we uncover an indirect role for microRNA expression in transgene silencing through effects on Polycomb complex expression. Additionally, we show that a commonly used marker gene, *Stubble* (*Sb*), affects Polycomb complex expression, which may be important in interpreting experiments assaying Polycomb function in *Drosophila* development. By ruling out a plausible candidate for TEI at the Fab2L transgene our work highlights the variability in different modes of TEI across species.

## 1. Introduction

In addition to the core genetic information encoded in DNA, eukaryotes possess additional layers of epigenetic information which affect gene expression without altering DNA sequence [1]. This epigenetic information includes signals such as DNA methylation [2], histone modifications [3] and small non-coding RNAs (sRNAs) [4], which form a key part of gene regulatory pathways in both development and adaptation, contributing to differences between cells within an individual as well as between individuals within a population. Often, multiple epigenetic mechanisms combine to produce robust epigenetic states [5].

In some cases, epigenetic information can be transmitted through the germline to subsequent generations in a process known as Transgenerational Epigenetic Inheritance (TEI) [6]. While the precise mechanism by which these epigenetic signals are inherited is often unclear, proposed mechanisms usually invoke either the direct “replicative” inheritance of the signal through the gametes, or a form of indirect, “reconstructive” inheritance whereby one signal is erased during development but reconstructed later from a different, directly inherited signal [7].

sRNAs have been implicated in many of the best described mechanisms of TEI [8–13]. In some organisms, they also have well-described mechanisms of gametic transmission [12,14,15] and self-propagation [16,17]. In *Drosophila melanogaster*, the major classes of sRNAs are 22-nucleotide (22nt) micro-RNAs (miRNAs), which recognise transcripts through short complementary base pairing target sites in the mRNA 3’-untranslated region [18], 21-24nt small interfering RNAs (siRNAs), which silence target transcripts via the RNA-induced silencing complex (RISC) [17,19], and 22-28nt piwi-interacting RNAs (piRNAs), responsible for both transcriptional and post-transcriptional silencing of transposable elements [19–21].

The mechanisms whereby sRNAs contribute to TEI are best described in *Caenorhabtitis elegans*. Key to this is the activity of RNA-dependent RNA polymerases which synthesise a type of sRNA known as 22G-RNAs. 22G-RNAs in association with Argonaute proteins, notably the nuclear Argonaute HRDE-1, can be transmitted through the germline and recruit RNA-dependent RNA polymerase, leading to synthesis of more 22G-RNAs and thus stable epigenetic memory. Initial targeting of 22G-RNAs is often kick-started by piRNA targeting, providing a mechanism whereby piRNAs can initiate stable silencing that can last even if piRNAs are removed [22–24].

*Drosophila melanogaster* does not have RNA-dependent RNA polymerase. Nevertheless, piRNA-mediated silencing of transposons in *Drosophila*, which operates partly through recruitment of the repressive histone modification H3K9me3, has also been shown to be transgenerationally inherited [25,26]. Some other cases of TEI in *Drosophila*, however, have reported the inheritance of histone modifications not usually associated with sRNA-directed silencing, in particular H3K27me3 [27–29]. The potential involvement of sRNAs in these cases has yet to be fully explored and is an important step in determining if these histone modifications can truly be inherited directly, rather than simply acting as secondary signals reconstructed from inherited sRNAs.

One such case is the transgenic *Drosophila* line Fab2L [30,31]. This line displays TEI involving either the silencing or activation of a transgenic region including a *mini-white* reporter gene, responsible for pigmentation in the adult eye. While eye colour is initially highly variable, these flies can be selected to produce genetically identical “epilines” with either fully white or fully red eyes [29]. Once established, these epilines can be maintained for many generations (>100) in the absence of selection. The primary epigenetic signal responsible for these phenotypic differences has been identified as the histone modification H3K27me3, deposited by the Polycomb Repressive Complex 2 (PRC2). However, whether sRNAs are also involved in this heritable epigenetic phenotype is not known.

In order to investigate the potential involvement of sRNAs in this case of TEI, we performed small RNA sequencing (sRNA-seq) in several different Fab2L lines as well as analyses of Fab2L lines bearing mutations in key sRNA pathway genes. We detected no evidence of sRNAs mapping to the Fab2L transgene, nor any effect of most sRNA mutations on eye colour, its inheritance or the expression of key genes involved in Fab2L TEI. We thus conclude that sRNAs do not play a role in the epigenetic phenotypes of the Fab2L line. However, we report a novel effect of the *Sb[1]* mutation on the expression of the PRC2 component *Pleiohomeotic* (*Pho*), which may have wider-ranging implications for the interpretation of results involving this widely-used genetic marker in other studies involving Polycomb targets.

## 2. Results and Discussion

### 2.1 Small RNAs do not contribute to phenotypic differences between genetically identical epilines

The *Drosophila* Fab2L line carries a single copy 12.4kb transgene inserted into chromosome 2 [29,30]. This transgene contains the reporter genes *LacZ* and *mini-white* under the control of the *Fab-7* regulatory element, which is also present endogenously on chromosome 3. The *mini-white* reporter gene, which controls red pigment deposition in the eye, is not expressed uniformly in Fab2L flies but shows a mosaic pattern of eye pigmentation, with some ommatidia showing strong *mini-white* expression and others strong repression (figure 1*a*). This variability is attributed to the stochastic binding of the Polycomb Repressive Complex 2 (PRC2) to the *Fab-7* element, which deposits the repressive histone modification H3K27me3 to randomly silence the transgene during development in some cells, but not in others [29,32].

**Figure 1.**
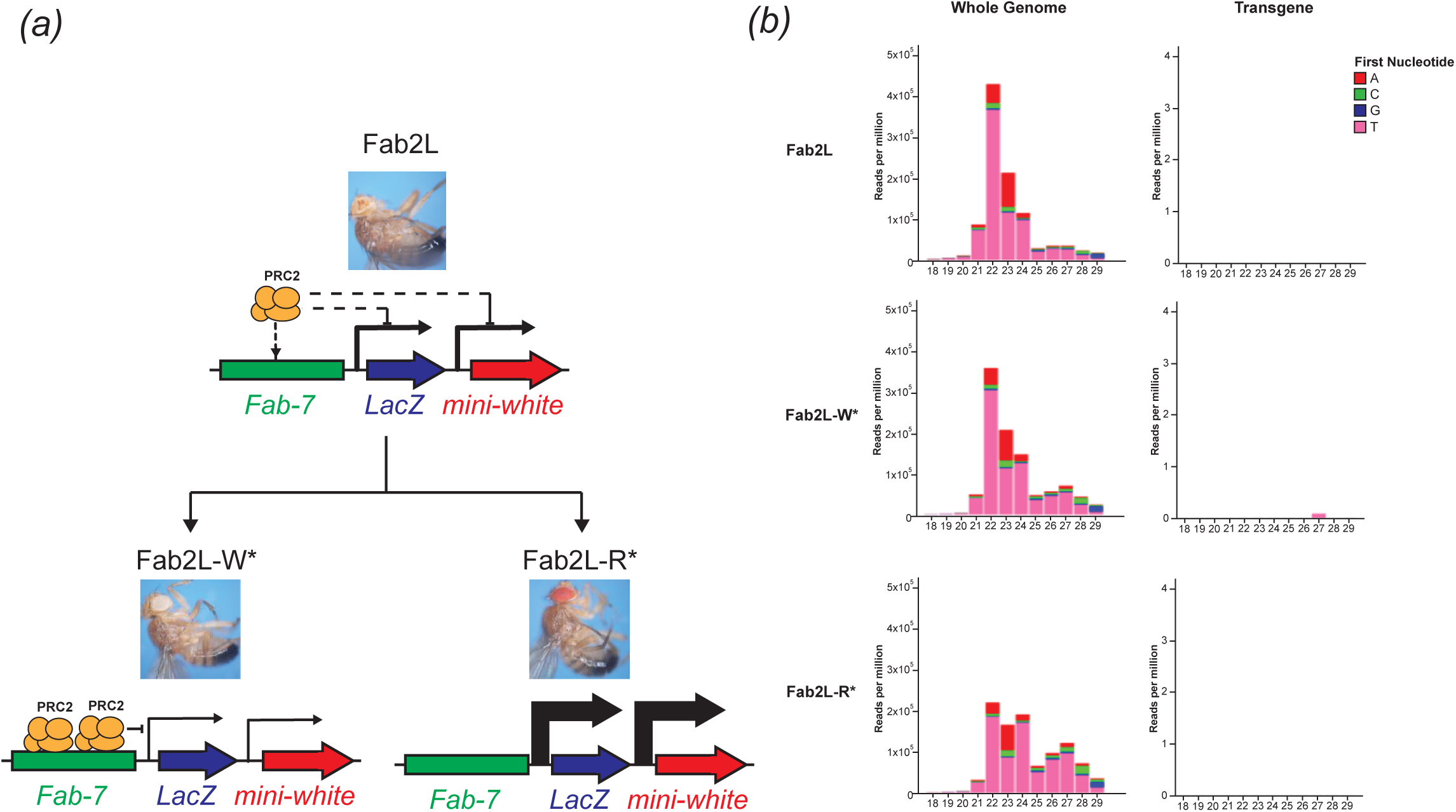
No small RNAs map to the Fab2L transgene in either naïve flies or epilines. ***(a)*** Shematic representation of the Fab2L transgene, containing *LacZ* and *mini-white* reporter genes under control of the *Fab-7* regulatory element. Stochastic binding of PRC2 to *Fab-7* leads to random inactivaton of the transgene, resulting in a mosaic eye colour which can be selected to produce either fully white or fully red-eyed fly lines. ***(b)*** sRNA-seq in embryos of the indicated genotypes. 18-29nt reads were mapped to either the whole *Drosophila* genome (left) or the Fab2L transgene sequence (right). This latter includes the *Fab-7* element with homology to the endogenous *Fab-7*, thus any reads mapping to the endogenous sequence would also map to the transgene. Colours indicate the first nucleotide of each read.

Transgenerational inheritance of these epigenetic differences over the transgene can lead to the establishment through artificial selection of genetically identical but phenotypically distinct “epilines”: populations of flies in which all individuals have either fully red or fully white eyes (figure 1*a*). Phenotypic differences between these epilines correlate with different levels of H3K27me3 across the transgene, both in the adult head and in the embryo [29]. However, to date no other epigenetic differences between the epilines have been identified, nor has a potential role for small RNAs been tested.

To determine if small RNAs contribute to the phenotypic differences between Fab2L-W* (white-eyed) and Fab2L-R* (red-eyed) epilines, we performed sRNA-seq in embryos from both of these epilines, as well as from unselected mosaic Fab2L flies. Despite good coverage of the genome as a whole, including many 21-28nt reads corresponding to the expected length of sRNAs, almost no sRNA transcripts mapped to either the transgene, or the endogenous *Fab-7* element in any of the lines (figure 1*b* and supplementary material, figure S1). This indicates that there is no direct targeting of the Fab2L locus or its transcripts by sRNAs. Given that all previous examples of sRNA mediated epigenetic inheritance involved direct targeting of the locus by sRNAs [33,34], the absence of sRNAs indicated that they were unlikely to contribute to the heritable epigenetic differences in eye colour between these populations.

### 2.2 Initiation of epigenetic inheritance through chromatin contacts does not involve small RNAs

We next sought to determine whether small RNAs might contribute to the initiation of epigenetic inheritance. Epigenetic inheritance of eye colour in Fab2L does not happen spontaneously, but requires a triggering event. Indeed, in a ‘naïve’ Fab2L population, eye colour is not heritable (Figure 2*a*). Epigenetic inheritance can be established by introducing a single generation of heterozygosity at the endogenous *Fab-7* locus [29]. After this transient genetic change, the resultant Fab2L flies, although genetically identical to the naïve Fab2L line, now have heritable eye colour and can be selected to produce epilines with either red or white eyes (Figure 2*b*). Previous work has shown that establishment of TEI involves chromatin contacts between the transgenic and endogenous *Fab-7* elements, which are present in Fab2L flies and increase even further upon *Fab-7* heterozygosity. These contacts alone are sufficient to trigger TEI, but are not required for the maintenance of heritable eye colour differences [32]. Furthermore, sRNA pathways were previously implicated in contacts between *Fab-7* elements in a related fly line carrying an X-chromosome transgene [35]. We therefore asked whether small RNAs could similarly be contributing to the initiation, rather than the maintenance, of heritable epigenetic changes.

**Figure 2.**
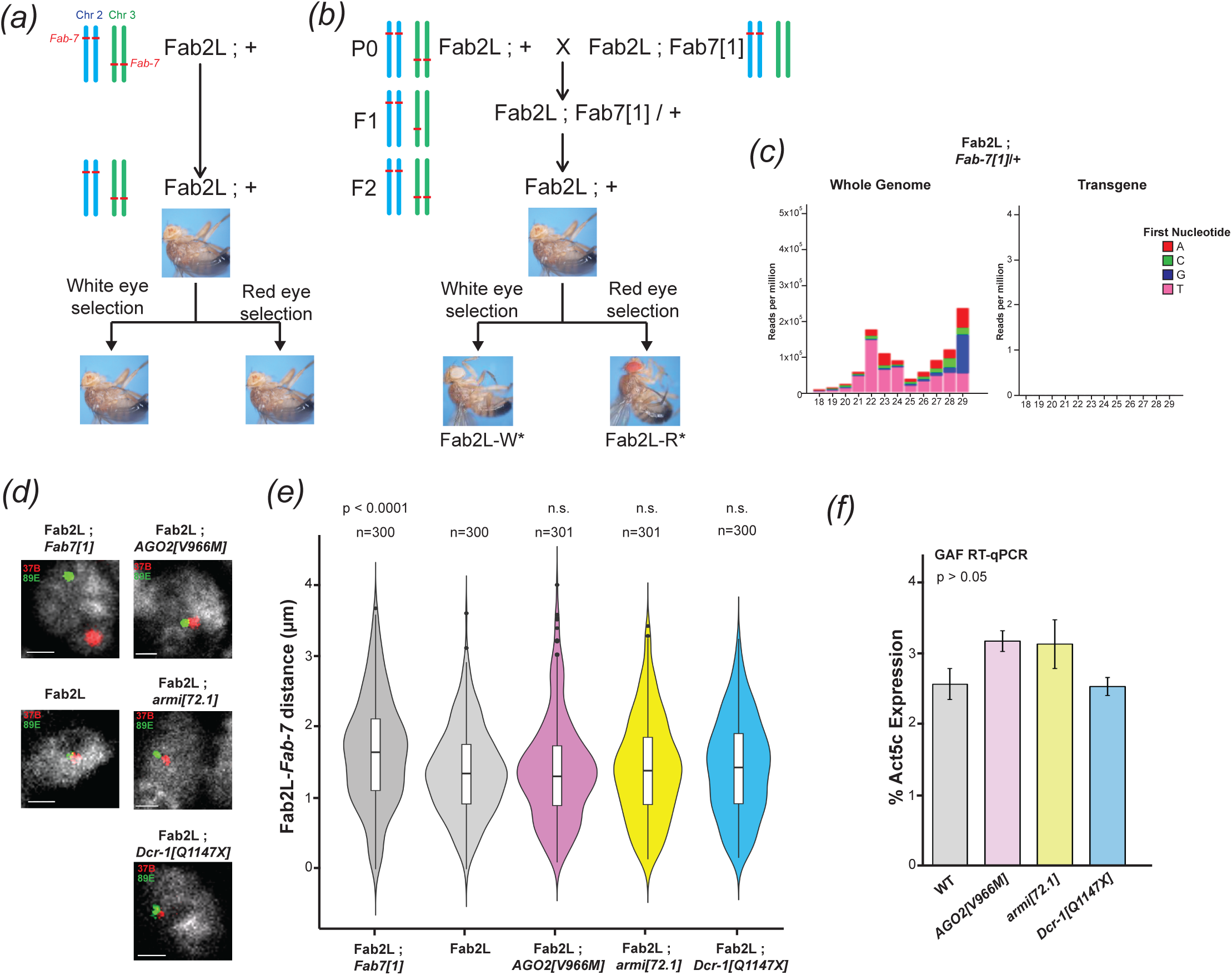
Small RNA pathways do not contribute to TEI establishment either in concert with or independently from chromatin contacts. ***(a,b)*** Crossing scheme for the triggering of TEI at Fab2L, with diagrammatic representation of the copy number of the *Fab-7* element on chromosomes 2 and 3. Heterozygosity of the endogenous *Fab-7* on chromosome 3 leads to establishment of TEI at the transgenic site on chromosome 2, thus enabling selection of epilines. ***(c)*** sRNA-seq in Fab2L ; *Fab7*[1]/+ embryos heterozygous for the endogenous *Fab-7*. 18-29nt reads were mapped to either the whole *Drosophila* genome (left) or the Fab2L transgene sequence (right). Colours indicate the first nucleotide of each read. ***(d)*** Illustrative micrographs of FISH in embryonic nuclei of the indicated genotypes. Nuclei are stained with DAPI, the 37B locus surrounding he Fab2L transgene is stained in red the 89E locus surrounding the endogenous *Fab-7* is stained in green. Scale bars represent 1µm. ***(e)*** Violin plots representing the distribution of average distance between the 37B and 89E regions surrounding the Fab2L transgene and endogenous *Fab-7*, respectively, as determined by FISH in the indicated genotypes. Distance were measured in stage 14-15 embryos in T1 and T2 segments. Significance was determined by two-way ANOVA, with distributions compared using Tukey’s HSD (n.s. = p > 0.1). ***(f)*** RT-qPCR for expression of GAF in Fab2L embryos homozygous for the indicated mutations. Expression was normalized to Act5C and overall significance was tested by one-way ANOVA.

To test this, we performed sRNA-seq on Fab2L ; *Fab7*[1]/+ embryos heterozygous for the endogenous *Fab-7* and corresponding to the time period when epigenetic inheritance is initiated (Figure 2*c*, and supplementary material, figure S2). We did not detect small RNA transcripts mapping to either the transgene or the endogenous *Fab-7*, arguing against the involvement of small RNAs in the initiation of TEI.

To test these conclusions further, we examined the effects of mutations that disrupt small RNA pathways on the Fab2L-*Fab7* chromatin contacts which are essential for TEI initiation. We performed fluorescence in situ hybridisation (FISH) on embryos using probes that highlight the regions surrounding the *Fab-7* elements (Figure 2*d*). As previously reported, these two regions frequently colocalize in the nuclei of Fab2L embryos, but not in Fab2L ; *Fab7*[1] embryos which lack the endogenous *Fab-7*, resulting in a significant decrease in the average distance between the two loci measured across many nuclei (Figure 2*e*, p < 5e-6, Tukey’s HSD after two-way ANOVA). We then looked at the frequency of contacts in Fab2L embryos carrying homozygous loss-of-function mutations in three genes with major roles in each of the three small RNA pathways. These were *Argonaute 2* (*AGO2*, required for siRNA-directed silencing [36]), *armitage* (*armi*, required for piRNA biogenesis [37–39]) and *Dicer-1* (*Dcr-1*, required for miRNA biogenesis [40]). All three mutant embryos showed a similar frequency of contacts to the wild-type Fab2L embryos (Figure 2*d,e*). Accordingly, we also found no effect of any of these mutations on expression of the gene *Trithorax-like* (*Trl*), also known as *GAGA-Factor* (*GAF*), which was previously shown to be the primary factor driving the contacts between the transgenic and endogenous *Fab-7* elements (Figure 2*f*).

Taken together, these results indicate that small RNAs are not involved in the initiation of TEI, either through direct effects on the transgene or indirectly by contributing to the previously observed increase in Fab2L-*Fab7* chromatin contacts. Both establishment and inheritance of the Polycomb-dependent epigenetic phenotypes in Fab2L therefore appear to be independent of sRNA pathways.

### 2.3 Horizontal transfer of epigenetic state by paramutation is independent of small RNAs

The epigenetic state of the Fab2L transgene can be transmitted horizontally between alleles by the process of “paramutation” [29]. Paramutation denotes a type of non-mendelian inheritance whereby an epigenetic state is transmitted *in trans* between two homologous alleles [41]. In the Fab2L line, crossing a naïve Fab2L with an established Fab2L epiline (either white or red-eyed) can result in the naïve allele acquiring the altered epigenetic state of the epiline allele. This phenomenon can be tracked by the use of a recessive *black[1]* marker allele, closely linked to the Fab2L transgene, such that F2 individuals that have inherited both copies of Fab2L from the naïve parent can be determined with high probability (supplementary material, figure S3). Although these F2 flies possess the genetic material of the naïve P0 population, the majority have an epigenetic state more closely resembling that of the epiline with which it was crossed, demonstrating that they have acquired a new epigenetic state.

Small RNAs have been implicated in paramutation in many organisms, including *Drosophila*, where they are often proposed to be the carrier of epigenetic information that transmits the epigenetic state from one allele to the other [33,42]. However, the mechanism of paramutation at the Fab2L locus remains unclear. In order to test the potential role of small RNA pathways in Fab2L paramutation, we performed crosses between Fab2L epilines and naïve Fab2L flies bearing different mutations to determine their effect on the efficiency of paramutation. Due to experimental constraints imposed by the location of the Fab2L transgene on chromosome 2, we tested only mutations in genes located on chromosome 3. In each case, we crossed both white and red epilines with a naïve Fab2L line that was both homozygous for the *black[1]* allele (closely linked to the Fab2L transgene) and heterozygous for a mutation balanced on the chromosome 3 balancer TM3-Ser (figure 3*a*). In the F1 generation, we sorted the flies into two populations, those that inherited the balancer and those that inherited the mutation, which were then self-crossed in parallel. The recessive *black[1]* phenotype then allowed us to select for F2 flies from each of these crosses that had inherited both copies of the Fab2L transgene from the naïve parent, which we then scored for eye colour and compared. Thus, the effect of each mutation on the F2 population could be compared to an internal control derived from the same initial P0 cross. This was important, as variations between the eye colour of starting populations can impact the F2 phenotype, making comparisons between crosses unreliable. Differences between these F2 populations would indicate an effect of the mutation on Fab2L eye colour, suggestive of a role for the mutated gene either in the horizontal transfer of epigenetic information by paramutation, or more directly in regulating expression of the transgene.

**Figure 3.**
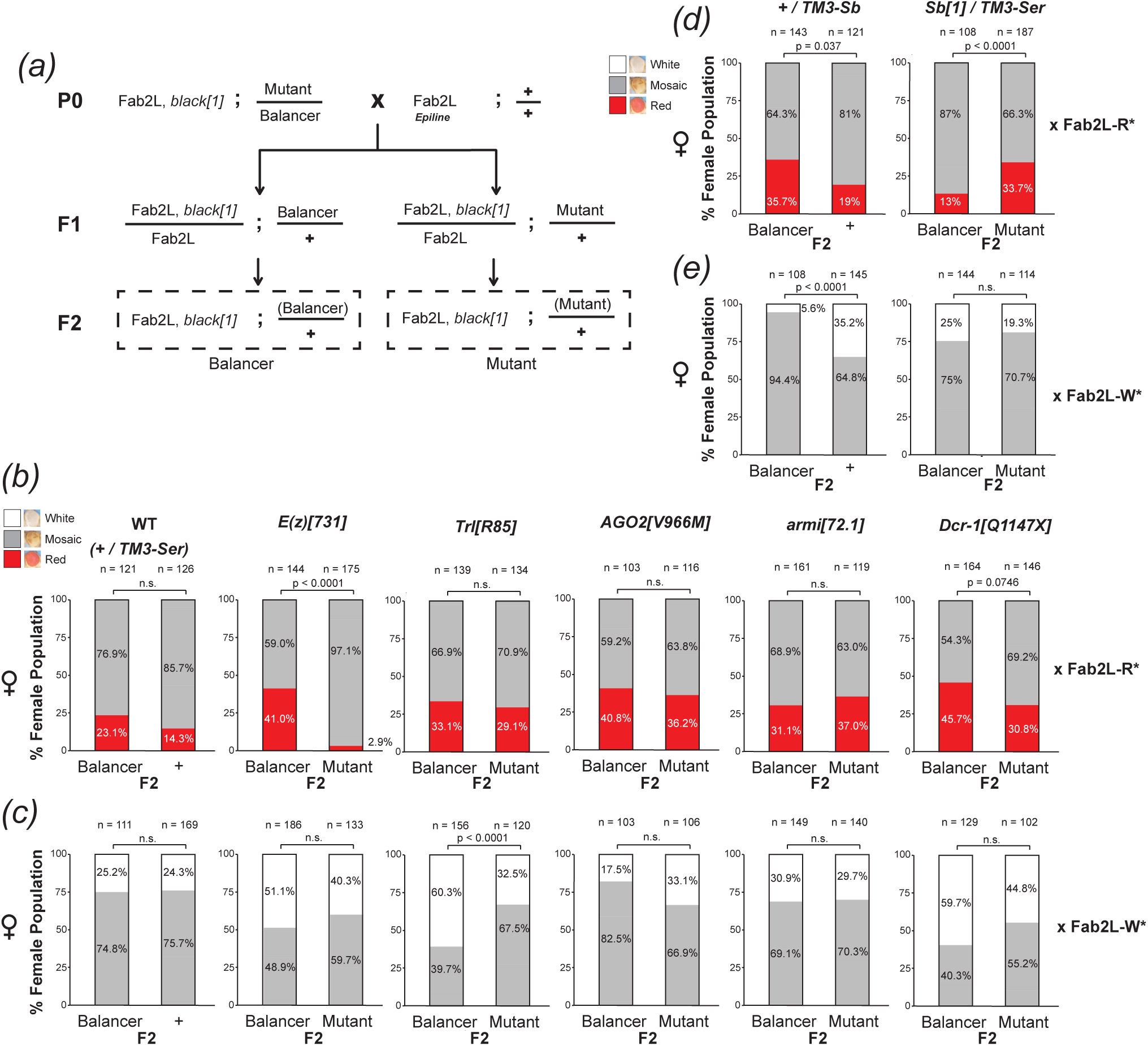
Mutation in *Dcr-1* and *Sb*, but not *AGO2* or *armi*, interfere with paramutation. ***(a)*** Crossing scheme to test the effect of mutations on the efficiency of paramutation. In each case naïve Fab2L flies carrying the transgene link to *black[1]* as well as a chromosome 3 mutation balanced on TM3-Ser were crossed with an estabished white or red epiline. F2 flies having inherited the transgene from the naïve P0 and the mutation were compared to those that had inherited the naïve transgene and the balancer as an internal control derived from the same cross. ***(b-e)*** Phenotypic distribution of eye colour in the F2 females of the paramutation crosses. In the P0, naïve flies carrying the indicated mutation were crossed with either a red epiline (*b,d*) or a white epiline (*c,e*). In each case F2 adults from the same cross inheriting either the balancer or the mutant were scored for eye colour and compared by Fisher’s exact test, with p values adjusted by Bonferroni correction (n.s. = p > 0.1).

We performed these crosses with naïve Fab2L flies bearing the previously mentioned loss-of-function mutations for *AGO2*, *armi* and *Dcr-1*. As controls, we performed the same cross with Fab2L flies bearing no mutation on chromosome 3 (WT) as well as flies mutated for the essential PRC2 component *Enhancer of zeste*, or *E(z)* [43], and the chromatin organiser and insulator *Trl / GAF* [44], both of which have important roles in regulating the Fab2L transgene [29,32]. As expected, crossing WT flies with epilines resulted in F2 flies that had at least partially adopted epiline identity, with a large proportion of the population having fully red or fully white eyes, rather than the near-100% mosaicism observed in a naïve population (figure 3*b,c*, supplementary material, figure S4*a,b*). More importantly, there was no significant difference between F2 flies that had inherited the balancer, and those that had not. Conversely, mutations in both *E(z)* and *Trl* significantly reduced the number of monochrome-eyed flies in at least one of the crosses, compared to flies from the same cross that inherited the balancer (p < 2e-18 and p < 7e-6 respectively, Fisher’s exact test), consistent with the known role of these genes in regulating transgene expression.

Several cases of paramutation have implicated siRNAs in plants [41,42] and piRNAs in metazoans, including *Drosophila* [22,24,25,45]. However neither *AGO2* nor *armi* mutations had any effect on the efficiency of paramutation, arguing against their role in this particular case (figure 3*b,c*, supplementary material, figure S4*a,b*). Consistent with this, two other mutants in piRNA genes we tested, *maelstrom* (*mael*) [46] and *argonaute 3* (*AGO3*) [47], also had no effect (supplementary material, figure S4*c,d*) and we detected no small RNA transcripts mapping to the transgene in the F1 of our paramutation crosses (supplementary material, figure S5). Interestingly, however, mutation of *Dcr-1* had a small but measurable effect, most notably in the cross with the red epiline (figure 3*b*). While this difference was no longer statistically significant after adjustment by Bonferroni correction (p = 0.0075 before correction, p = 0.075 after correction, Fisher’s exact test), it remained within reportable range (p < 0.1) and represented a significant enough effect from a heterozygous mutation to merit further investigation.

### 2.4 The common genetic marker *Stubble* influences Fab2L transgene expression

When initially testing lines containing balancer chromosomes for use as controls, we performed the paramutation cross (figure 3*a*) using both Fab2L, *black[1]* ; + / TM3-Ser, bearing the scalloped wing marker due to mutation in the gene *Serrate* [48,49] and Fab2L, *black[1]* ; + / TM3-Sb, bearing the shortened bristle phenotype due to the *Sb[1]* mutation in the gene *Stubble* [48,50]. While the former displayed the expected similarities between the F2 populations, leading us to use TM3-Ser in all subsequent crosses (figure 3*b,c*), the TM3-Sb containing line did not. Indeed, F2 flies that inherited the TM3-Sb balancer showed a significant shift towards red eyes compared to those that did not (p = 0.037 for red cross, p < 7e-9 for white cross, Fisher’s exact test) (figure 3*d,e* and supplementary material, figure S4*e*). To confirm that this difference reflected an effect of the mutation, we performed paramutation crosses with Fab2L flies carrying the *Sb[1]* mutation balanced over TM3-Ser. Once again, F2 flies that inherited the *Sb[1]* mutation showed a significant shift towards red eye colour colour (p < 7e-4 for red cross, Fisher’s exact test) (figure 3*d,e* and supplementary material, figure S4*e*). This unexpected result suggested that *Stubble* influenced the regulation of transgene expression in the Fab2L line.

### 2.5 *Dcr-1* and *Sb* mutations interfere with Polycomb recruitment by downregulating *Pho*

Though we set out to investigate the potential role of sRNAs in TEI at the Fab2L locus, we identified no discernible involvement of the two sRNA pathways most likely to be responsible for heritable epigenetic differences, siRNAs and piRNAs. Instead, we identified potential effects of the miRNA pathway (through mutation of *Dcr-1*) and *Stubble* on the epigenetic eye colour phenotype.

Unlike siRNAs and piRNAs, miRNAs have no method of self-propagation. They are however major post-transcriptional regulators of a wide array of genes. We reasoned therefore that the observed effect of *Dcr-1* mutation on Fab2L eye colour was unlikely to reflect an involvement in the inheritance of eye colour, but was more likely a secondary effect brought about by misregulation of one of the many miRNA targets. Similarly, *Stubble* is an important signalling protein whose mutation has many downstream effects. Its most clearly defined function is in epithelial morphogenesis, bringing about the visible bristle phenotype frequently used as a marker [50]. However, this function extends more generally to imaginal disc morphogenesis, with known phenotypes in both the leg and wing among others [51–55]. Its role in activating the Rho-GTPase signalling pathway [50] has also linked it to regulation of the Ecdysone receptor [53], a major transcription factor with hundreds of targets across developmental stages [56]. This led us to consider the possibility that mutation of *Sb* might affect transgene expression.

All mutations tested in our previous assays were homozygous lethal, disrupting as they do important genes with essential functions. Our analyses of adult eye colour were therefore limited to determining the effects of heterozygous mutants, which may mask some of the more severe effects of the mutations. All of these mutants, however, survive until at least late embryogenesis, if not the larval stage, allowing us to analyse homozygous mutants at earlier stages of development. To further investigate the effects of *Dcr-1* and *Sb* mutations, we performed RT-qPCR on Fab2L embryos homozygous for the previously used *Dcr-1* and *Sb* loss-of-function mutations. As controls, we used a naïve Fab2L line, a wild type w-line, containing no Fab2L transgene or endogenous *white* gene, and a line carrying a previously described mutant version of the Fab2L transgene (labelled here “Fab2L-constit.”) in which the binding sites for the PRC2 recruiters *Pho* and *GAF* are mutated, resulting in constitutive expression of the *mini-white* reporter in the transgene due to absence of PRC2 silencing [32]. We also analysed the *AGO2* and *armi* mutants which we had determined to have no effect on the transgene based on our paramutation crosses (figure 3*b,c*) to determine if homozygosity of these mutants revealed any affects not visible in heterozygotes.

The line carrying the mutated version of Fab2L showed strong expression of both *mini-white* and *LacZ* from the transgene, compared to the much lower but still detectable levels of expression in the unmutated Fab2L (figure 4*a,b*). Mutation of *AGO2* or *armi* had no significant effect on expression of either gene, confirming that the siRNA and piRNA pathways are not involved in regulation of the Fab2L transgene. However, mutation of both *Dcr-1* and *Sb* resulted in a significant increase in expression of *mini-white* and *LacZ* (p = 0.046 and p = 0.0046 for *Dcr-1*, p < 2e-5 and p = 0.069 for *Sb*, Tukey’s HSD test after one-way ANOVA). We observed upregulation of both transgenic reporter genes, which are the products of two different transcripts whose expression is driven by the same promoter, thus indicating that the effects were at the level of transcription, rather than involving post-transcriptional regulation of *mini-white* by Dcr-1, or potential further downstream effects on eye morphogenesis by Sb. We therefore concluded that both Sb and Dcr-1 were likely to regulate transcription of the transgene indirectly.

**Figure 4.**
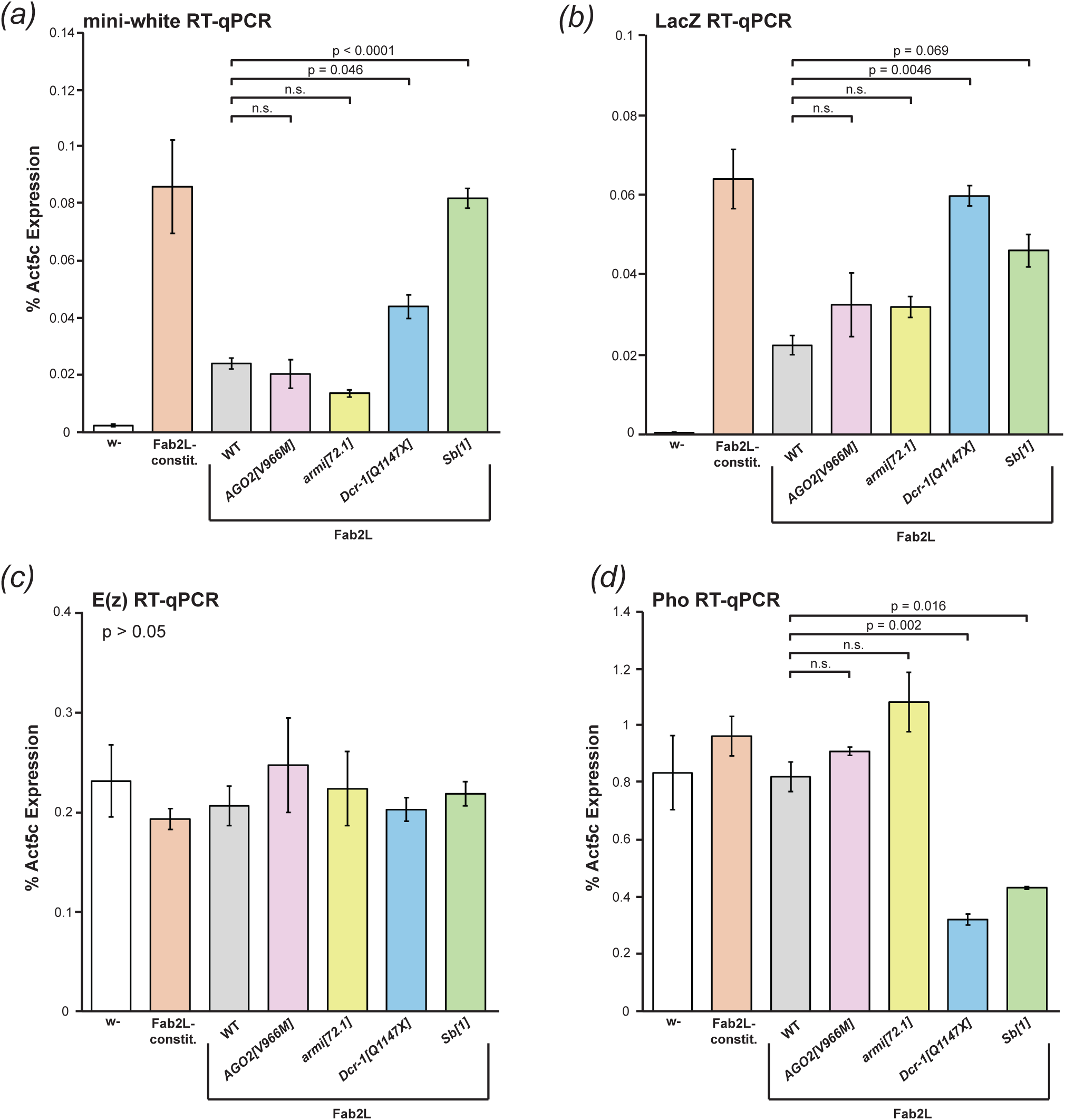
Mutation in *Dcr-1* and *Sb* increase transcription from the transgene and decrease expression of Pho. ***(a-d)*** RT-qPCR for expression of mini-white (*a*), LacZ (*b*), E(z) (*c*) and Pho (*d*) in embryos of the indicated genotypes. Expression was normalized to Act5C. Overall significance in the Fab2L samples was tested by one-way ANOVA, followed by pair-wise comparison using Tukey’s HSD in the event of significance (n.s. = p > 0.1).

Transcriptional regulation of the transgene is primarily controlled by PRC2, which deposits H3K27me3 to silence the transgene to varying degrees in the naïve, white and red epilines of Fab2L [29,32]. To determine if *Dcr-1* and *Sb* mutation were affecting the activity of PRC2, we performed RT-qPCR to measure the PRC2 catalytic subunit *E(z)*, responsible for depositing H3K27me3 [57], and *Pho*, which recruits PRC2 to its targets by recognizing specific binding sites, including several within the *Fab-7* element [32,58]. *E(z)* expression was similar across all lines tested (figure 4*c*), whereas *Pho* was significantly downregulated in both the *Dcr-1* and *Sb* mutant lines (p = 0.002 and p = 0.016, respectively, Tukey’s HSD after one-way ANOVA) (figure 4*c*). This suggested a potential mechanism by which both *Dcr-1* and *Sb* mutations affect Fab2L eye colour, downregulating *Pho* and interfering with the binding of PRC2 to the *Fab-7* element within the transgene. Exactly how these effects on *Pho* expression are brought about is difficult to determine, but the involvement of miRNAs in many gene expression networks suggests the possibility that loss of Dcr-1 could lead to alterations in regulation of *Pho*.

To our knowledge this effect of Sb on *Pho* has not previously been reported. This may be explained by the fact that the decrease in *Pho* expression is modest (around 2-fold). Its effects may not manifest in other developmental phenotypes due to redundancy and reinforcement in endogenous gene expression networks providing a buffer to potential environmental or genetic perturbations [59]. Polycomb targets in particular are often coregulated in a manner that may be dependent on genomic location and 3D chromatin organisation [60]. As a transgene carrying a Polycomb-bound regulatory sequence divorced from its natural context, Fab2L may be more susceptible than endogenous loci to perturbation from mild disruptions that would normally be absorbed by these safeguards. Heterozygosity of *Pho*, for instance, is unlikely to have a significant effect on most Polycomb targets due not only to the presence of the remaining wild-type *Pho* copy, but also redundancy with the related *Pho-like* and other PRC2 recruiters and coregulation with other targets [57,58,61]. Eye colour in Fab2L, on the other hand, reflects a stochastic pattern established by Pho-directed PRC2 binding in early development, and maintained through to adulthood. This fragile system is thus much more susceptible to disruption. Moreover, the possibility of observing the effect across many individuals, each with a large number of ommatidia in their eyes, is more likely to reveal what in essence remains a mild effect on the average phenotype of the population (figure 3*d,e*).

Nevertheless, the ubiquity of *Sb[1]* as a marker may warrant caution in its use in certain cases. Indeed, the mild effects of a mutation may be exacerbated when combined with additional stressors such as extreme temperatures, something to which Polycomb targets may be particularly susceptible [62,63]. It may therefore be prudent to keep these effects in mind for future analyses involving Polycomb-regulated genes balanced on the TM3-Sb balancer, or otherwise linked to the *Sb[1]* mutation, and perhaps favour the use of the alternative TM3-Ser or other markers in these cases.

### 2.6 Conclusion

In this study we investigated the potential involvement of sRNAs in the transgenerational inheritance of a Polycomb-dependent epigenetic eye colour phenotype in *Drosophila melanogaster*. Both sequencing and mutational analysis found no evidence for the involvement of either siRNAs or piRNAs in the establishment or maintenance of heritable epigenetic differences between individuals. These results provide further evidence that histone modifications, in this particular case H3K27me3, can be inherited transgenerationally in the absence of any other associated mark, encouraging further work to elucidate the exact mechanism by which this inheritance occurs. We identified indirect effects of the miRNA pathway on the expression of PRC2 component *Pho*, with interesting implications for understanding the control of PRC2-mediated regulation during development. Moreover, we also discovered an effect on *Pho* of the *Sb[1]* mutation, highlighting a previously unknown interaction between this marker gene and the Polycomb silencing pathway.

## 3. Material and methods

### 3.1 Fly stocks and culture

Flies were raised in standard cornmeal yeast extract media. Standard temperature was 21°C, with the exception of egg laying for RNA extraction which was performed at 18°C. The Fab2L and Fab2L ; *Fab7[1]* lines were described in Bantignies *et al.*, 2003[30]. The Fab2L, *black[1]* line and pre-established Fab2L epilines (Fab2L-R* and Fab2L-W*) were described in Ciabrelli *et al.*, 2017[29] while the Fab2L-constit. (also called Fab2L-INS-PRE) was described in Fitz-James et al. 2023[32].

Fab2L, *black[1]* ; *Trl^R85^*/TM6, Fab2L, *black[1]* ; *E(z)^731^*/TM3-sb and Fab2L, *black[1]* ; *Sb[1]*/TM3-Ser flies were described in Ciabrelli *et al.*, 2017[29] and were crossed together to generate Fab2L, *black[1]* ; *Trl^R85^*/TM3-Ser, Fab2L, *black[1]* ; *E(z)^731^*/TM3-Ser, Fab2L, *black[1]* ; *+*/TM3-Sb and Fab2L, *black[1]* ; *+*/TM3-Ser.

Lines bearing the mutations *AGO2[V966M]* (BL32062), *armi[72.1]* (BL8544), *Dcr-1[Q1147X]* (BL32066), *mael [r20]* (BL8516) and *AGO3[t3]* (BL28270) were ordered from the Bloomington Drosophila Stock Centre and crossed with Fab2L, *black[1]* ; *Sb[1]*/TM3-Ser to generate the balanced lines.

To obtain homozygous mutant embryos, the above mutant lines were balanced on the TM3, Sb, Kr-GFP (TKG) balancer chromosome, described in Casso et al. 1999[64], by crossing them with *Pc*/TKG flies gifted by the Cavalli lab. The TKG balancer contains a fluorescent GFP marker expressed under control of the *krüppel* promoter, giving a distinct GFP pattern in mid to late embryos, thus allowing for selection of homozygous embryos (see below).

### 3.2 Embryo Collection

For both RNA extraction and FISH, flies were put in cages with yeasted apple juice agar plates for egg-laying. Embryos were collected and washed in dH2O before dechorionation with bleach. Embryos were collected either at stage 12-13 to allow time for sorting, or stage 14-15 if proceeding directly to RNA extraction or FISH. Homozygous mutant embryos were obtained by sorting for GFP negative (or very strong GFP expression in the case of *Sb* mutants) embryos after dechorionation and kept in dH2O or Buffer A until ready. For RNA extraction, some embryos were stored at −20°C prior to extraction.

### 3.2 RNA extraction

A minimum of 20 embryos per sample were washed in DEPC-treated H2O and pelleted with gentle centrifugation at 4,000xg for 2 minutes. Embryos were then homogenized in 50μl Trizol reagent using a disposable pestle and then incubated at room temperature for 5 minutes in a total volume of 1mL Trizol. Samples were mixed with 200μl Chloroform, incubated 2 minutes at room temperature and centrifuged 12,000xg for 15 minutes at 4°C. The aqueous supernatant was taken and RNA precipitated with 500μl isopropanol for 10 minutes at 4°C. Samples were centrifuged 12,000xg for 10 minutes, washed in 1mL 75% EtOH and centrifuged again. The pellet was air dried and resuspended in 25μl RNase-free H2O. Residual DNA was removed using the DNA-free DNA Removal kit (Ambion) and RNA concentration determined using a Qubit spectrophotometer.

### 3.3 sRNA-seq

sRNA library preparation was performed by Novogene as was sequencing on an Illumina Novaseq 6000 using 50bp single-end reads. Starting from raw fastq files, adapters were trimmed using fastx_toolkit and sequences were aligned to the *Drosophila* genome (dm6) and a custom genome built from the Fab2L fasta sequence using Bowtie-build, using 0 mismatches and reporting one alignment per read. Alignment files in sam format were converted to bam files using samtools and bam files were converted to bed files using bamToBed. The resulting bed files were then used as input for R scripts to collate the length and first nucleotide to produce plots. R Code to reproduce the plots is available via the SarkiesLab github page (SarkiesLab/Fab2LsRNA).

### 3.4 RT-qPCR

200ng RNA was converted to cDNA using the High-Capacity RNA-to-cDNA kit (Applied Biosystems) before dilution 1:40. Samples were then subjected to qPCR in triplicate on a CFX 384 Real-time PCR System (Bio-Rad) with a total volume of 10μl per well (5 μl cDNA and 5 μl iTaq universal SYBR green supermix containing 1μM of each primer). Cq values were averaged across the three triplicates and normalized to Act5C.

### 3.5 Fluorescence in situ hybridization

Two-color 3D FISH was performed as previously described[65]. For a detailed protocol, see Bantignies and Cavalli, 2014[66]. Embryos collected as indicated above were fixed at stage 14-15 in buffer A (60 mM KCl; 15 mM NaCl; 0.5 mM spermidine; 0.15 mM spermine; 2 mM EDTA; 0.5 mM EGTA; 15 mM PIPES, pH 7.4) with 4% paraformaldehyde for 25 min in the presence of heptane then devitellinized by adding methanol to the heptane phase, extracted and washed three times in methanol. Embryos were kept for a maximum of 4 months in methanol at 4C before proceeding to FISH. Fixed embryos were sequentially re-hydrated in PBT (PBS, 0.1% Tween 20) before being treated with 100–200 μg/ml RNaseA in PBT for 2 hours at room temperature. Embryos were then sequentially transferred into a pre-Hybridization Mixture (pHM: 50% formamide; 4XSSC; 100 mM NaH2PO4, pH 7.0; 0.1% Tween 20). Embryonic DNA was denatured in pHM at 80°C for 15 minutes. The pHM was removed, and denatured probes diluted in the FISH Hybridization Buffer (FHB: 10% dextransulfat; 50% deionized formamide; 2XSSC; 0.5 mg/ml Salmon Sperm DNA) were added to the tissues without prior cooling. Hybridization was performed at 37°C overnight with gentle agitation. Post-hybridization washes were performed, starting with 50% formamide, 2XSSC, 0.3% CHAPS and sequentially returning to PBT. After an additional wash in PBS-Tr, DNA was counterstained with DAPI (at a final concentration of 0.1 ng/μl) in PBT and embryos were mounted with ProLong Gold Antifade (Invitrogen).

FISH probes for the 37B and 89E regions were made from a previous design described in Ciabrelli *et al.* 2017[29]. For each region, 6 non-overlapping probes of between 1.2 and 1.7kb covering an area of approximately 12kb were generated using the FISH Tag DNA kit with Alexa Fluor 555 or Alexa Fluor 647 dyes (Invitrogen Life Technologies). 100ng of each probe were added to the 30µL of FHB for hybridization.

### 3.6 Microscopy and image analysis

For the FISH, the 3D distances between 37B and 89E loci were acquired and measured as follows: due to somatic pairing of homologous chromosomes in *Drosophila*, the majority of the nuclei in embryos show a single FISH spot for each probe. In the cases of non-overlap FISH signals between homologues, the closest distance between the centres of the two probes was considered. To measure distances, 3D stacks were collected from 3 different embryos. Optical sections were collected at 0.3-0.5 μm intervals along the Z-axis using a Zeiss LSM980 microscope at the Micron Imaging facility. Relative 3D distances between FISH signals were measured in approximately 100 nuclei per 3D stack using the Imaris software (Oxford Instruments) and plotted in R using the ggplot2 package.

### 3.7 Statistical analysis

Statistical analysis was performed in R using the Tidyverse package. Fab2L-*Fab7* distance measurements from the same embryo were assigned an embryo ID and distributions were tested by two-way ANOVA with the genotype and embryo ID as parameters, before pairwise comparison using Tukey’s HSD. RT-qPCR data was tested by one-way ANOVA and, in the case of significance, compared pairwise with Tukey’s HSD. Fisher’s exact test was used to compare the distribution of eye colour between the F2 of paramutation crosses. P-values were adjusted by Bonferroni correction. Given the correction applied to all p values, 0.1 was taken as the upper bound for significance, with all values below being reported.

## Supporting information

Raw Data File

## Ethics

This work did not require ethical approval from a human subject or animal welfare committee.

## Data accessibility

Access to primary data is available upon reasonable request to the corresponding authors.

## Declaration of AI use

We have not used AI-assisted technologies in creating this article.

## Author contributions

M. F-J. and P. Sarkies conceived of and led the project. M.F-J. and P. Sarkies designed the experiments. M. F-J. performed the sRNA-seq with P. Sparrow and the RT-qPCR and FISH with C. P. P. Sparrow performed the paramutation crosses and analysis. P. Sarkies analysed the sequencing data. M. F-J. and P. Sarkies interpreted the data. M. F-J. composed the manuscript with editorial input from P. Sarkies. All authors reviewed and commented on the manuscript.

## Conflict of interest delcaration

We declare we have no competing interests.

## Acknowledgements

We thank the Cavalli lab (Institute of Human Genetics, University of Montpellier and CNRS) for the gift of the Drosophila lines used in this study. We also thank the Micron Imaging Facility and the Jansen lab (Department of Biochemistry, University of Oxford) for the use of equipment.

---

**Figure S1.**
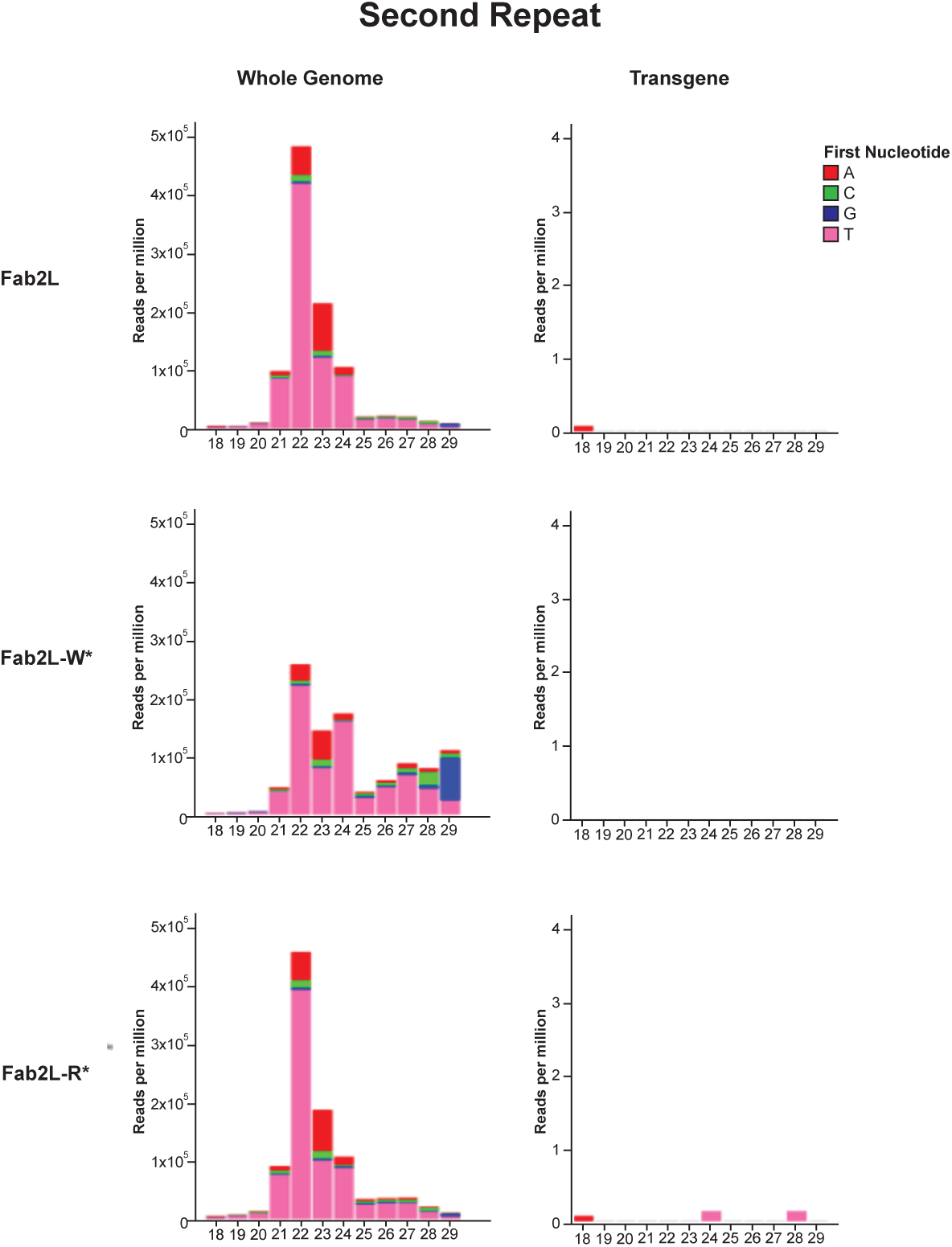
Additional sRNA-seq in Fab2L and epilines. Second repeat of the sRNA-seq of the indicated genotypes presented in figure 1*b*.

**Figure S2.**
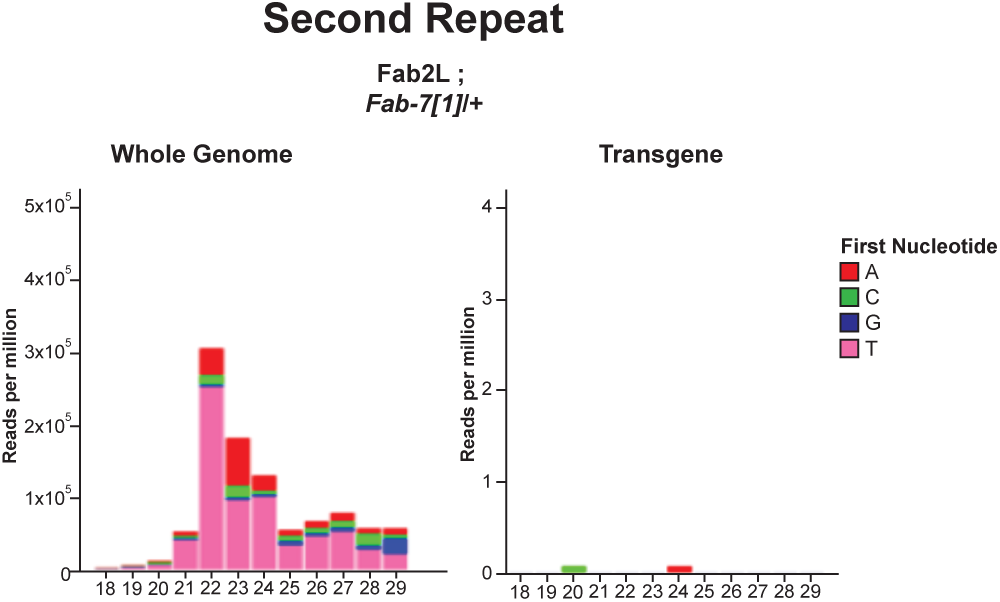
Additional sRNA-seq in Fab2L ; *Fab7[1]/*+. Second repeat of the sRNA-seq in Fab2L ; *Fab7[1]/*+ embryos presented in figure 2*c*.

**Figure S3.**
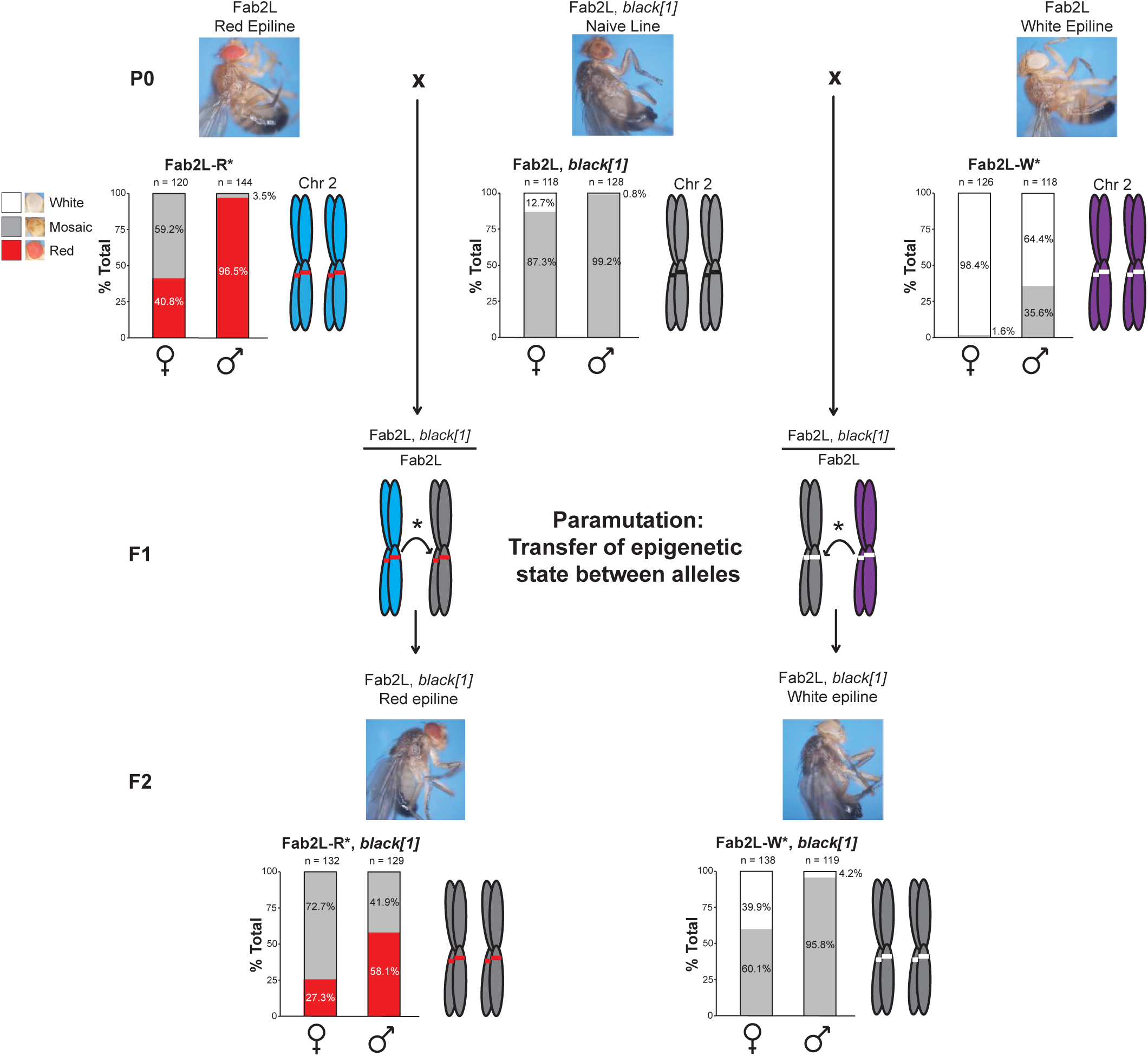
Paramutation allows the allelic transfer of epigenetic state between transgenes. Illustration of the basic paramutation crossing scheme. Naïve Fab2L flies carrying a homozygous *black[1]* mutation linked to the transgene (centre) are crossed to established epilines with high levels of fully red (left) or fully white (right) eyes within the population. In the F1, the altered epigenetic state of the epiline alleles (blue and purple chromosomes) are transferred *in trans* to the naïve alleles (grey chromosomes) by paramutation. Thus, F2 flies bearing two copies of the transgene inherited from the naïve P0 (as determined by the black phenotype) nonetheless have high levels of individuals with fully red or fully white eyes, attesting to the acquisition by these alleles of a new epigenetic state.

**Figure S4.**
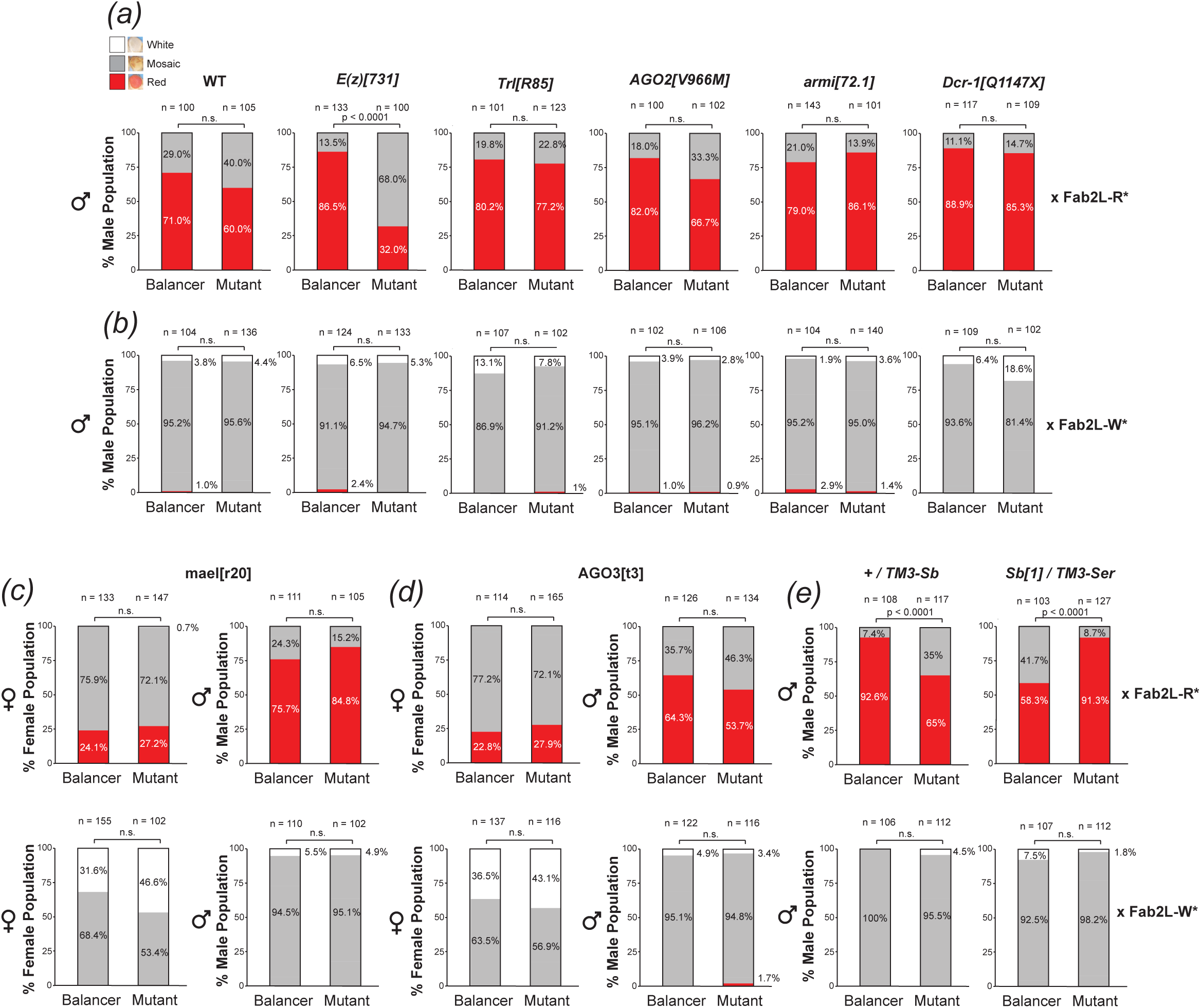
siRNA and piRNA mutants have no effect on paramutation efficiency. ***(a-e)*** Phenotypic distribution of eye colour in the F2 males of the paramutation crosses represented in figure 3 (*a,b,e*) and both females and males from additional crosses involving mutations in *mael* (*c*) and *AGO3* (*d*). In the P0, naïve flies carrying the indicated mutation were crossed with either a red epiline (*b,d*) or a white epiline (*c,e*). In each case F2 adults from the same cross inheriting either the balancer or the mutant were scored for eye colour and compared by Fisher’s exact test, with p values adjusted by Bonferroni correction (n.s. = p > 0.1).

**Figure S5.**
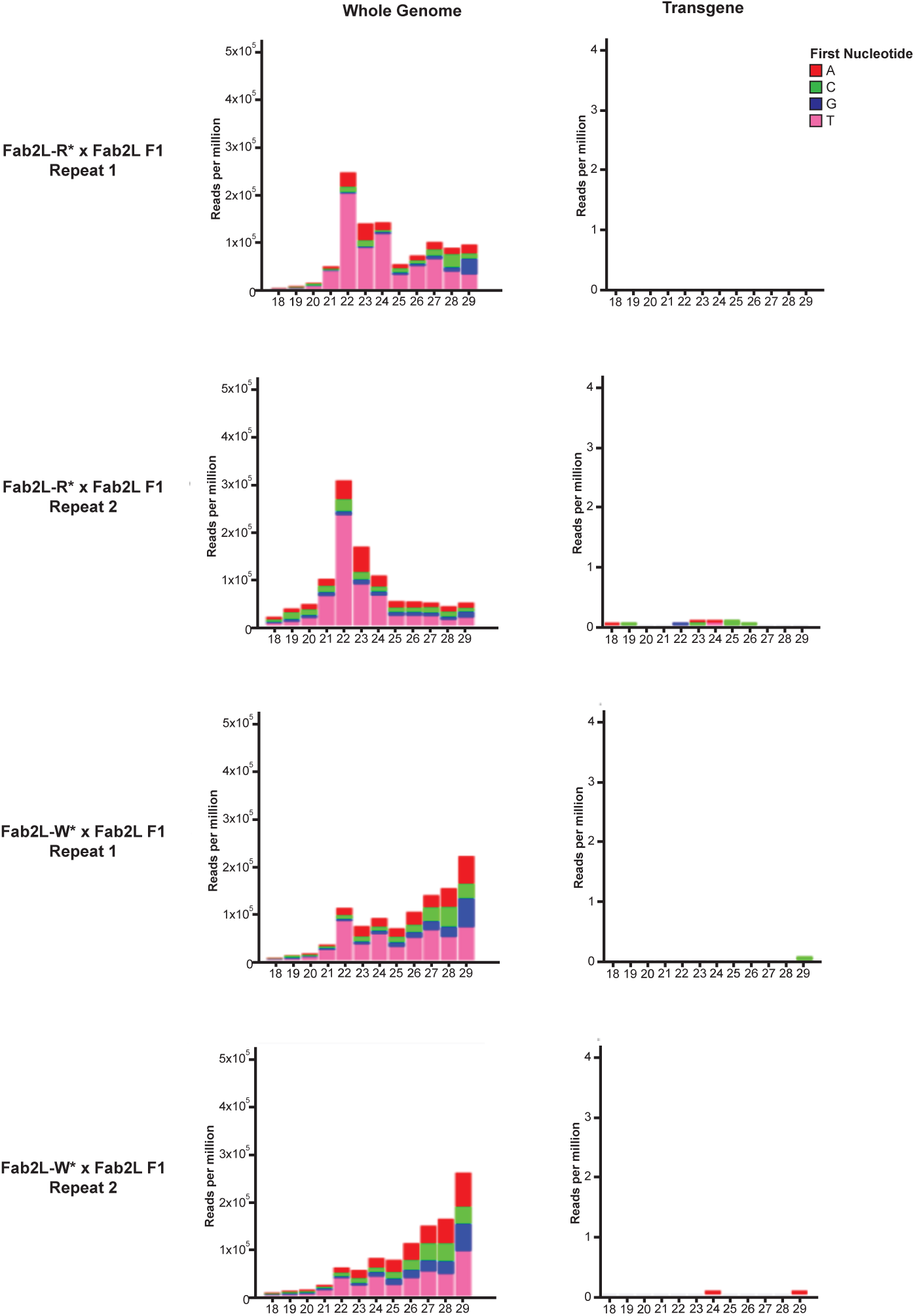
No sRNAs map to the transgene in the F1 of paramutation crosses. sRNA-seq in F1 embryos obtained by crossing naïve Fab2L with either red (top 4 panels) or white (bottom 4 panels) epilines. 18-29nt reads were mapped to either the whole *Drosophila* genome (left) or the Fab2L transgene sequence (right). Colours indicate the first nucleotide of each read.

